# The intracellular symbiont *Wolbachia* enhances recombination in a dose-dependent manner

**DOI:** 10.1101/686444

**Authors:** Kaeli N. Bryant, Irene L.G. Newton

## Abstract

*Wolbachia pipientis* is an intracellular alphaproteobacterium that infects 40-60% of insect species and is well known for host reproductive manipulations. Although *Wolbachia* are primarily maternally transmitted, evidence of horizontal transmission can be found in incongruent host-symbiont phylogenies and recent acquisitions of the same *Wolbachia* strain by distantly related species. Parasitoids and predator-prey interactions may indeed facilitate the transfer of *Wolbachia* between insect lineages but it is likely that *Wolbachia* are acquired via introgression in many cases. Many hypotheses exist as to explain *Wolbachia* prevalence and penetrance such as nutritional supplementation, protection from parasites, protection from viruses, or straight up reproductive parasitism. Using classical genetics we show that *Wolbachia* increase recombination in infected lineages across two genomic intervals. This increase in recombination is titer dependent as the *w*MelPop variant, which infects at higher load in *Drosophila melanogaster*, increases recombination 5% more than the *w*Mel variant. In addition, we also show that *Spiroplasma poulsonii*, the other bacterial intracellular symbiont of *Drosophila melanogaster*, does not induce an increase in recombination. Our results suggest that *Wolbachia* infection specifically alters host recombination landscape in a dose dependent manner.

**Article Summary:** The ubiquitous bacterial symbiont *Wolbachia* is known to alter host reproduction through manipulation of host cell biology, protect from pathogens, and supplement host nutrition. In this work we show that *Wolbachia* specifically increases host recombination in a dose dependent manner. Flies harboring *Wolbachia* exhibit elevated rates of recombination across the 2^nd^ and X chromosomes and this increase is proportional to their *Wolbachia* load. In contrast, another intracellular symbiont, *Spiroplasma*, does not lead to an increase in recombination across the intervals tested. Our results point to a specific effect of *Wolbachia* infection that may have a significant effect on infected insect populations.

## Introduction

Recombination, the exchange of genetic material during meiosis is thought to be largely beneficial, as it increases the efficacy of natural selection ^1,2^. Because of chromosome architecture, loci that are physically linked to each other can interfere with selection such that selection at one locus reduces the effective population size, and therefore the efficacy of selection, at linked loci. This phenomenon, termed “Hill-Robertson interference,” means that positive or negative selection at one site can interfere with selection at another site. By allowing loci to shuffle between chromosomes, recombination mitigates Hill-Robertson interference ^3^. As a result of this re-shuffling, areas of the genome subject to high recombination rates show higher nucleotide diversity, either because of the inherent mutagenic effect of recombination or by the indirect influence of recombination on natural selection in a population. Overall, a large body of literature supports the assertion that recombination increases efficacy of selection and enhances adaptation in animals, as studied in various *Drosophila* species ^1–3^.

One factor that may influence recombination is bacterial infection. For example, injection of flies with the bacterial pathogen *Serratia* increases recombination post infection ^4^. Many *Drosophila* species are colonized persistently by *Wolbachia pipientis*, an alpha-proteobacterium within the *Rickettsiales* and the most common infection on the planet, found in 40-60% of all insects. *Wolbachia’s* prevalence in populations is likely modulated by its reproductive manipulations, induced to benefit infected females ^5^. However, this reproductive parasitism alone is not sufficient to explain *Wolbachia* infection prevalence; indeed there are many recently discovered, insect infecting strains which do not seem to induce any reproductive phenotype at all, suggestive of other potential benefits provided by the symbiont ^6–8^. One known benefit is pathogen blocking, where *Wolbachia* repress the virus replication within the insect host ^9–11^. This phenomenon has important implications for the use of *Wolbachia* in vector control ^12^. In addition to protecting its host from pathogens, *Wolbachia* also generally improves the fitness and fecundity of some hosts, and removal of the endosymbiont can cause a decrease in host fitness (Fry and Rand ’02; Fry et al. ‘04). Finally, *Wolbachia* can rescue oogenesis defects in mutant *Drosophila* strains ^13^ and has also made itself a necessary component of oogenesis in some wasp species, thereby making the infection indispensable ^14^.

One recently discovered phenotype of *Wolbachia* is that it may increase the amount of recombination events on the X chromosome, but not on the 3^rd^, in *Drosophila melanogaster* ^15,16^. This phenotype contrasts with that observed for *Serratia* infection, where elevated recombination was observed on the 3^rd^ chromosome ^4^. Does *Wolbachia* infection actually lead to increased recombination? If so, would any infection of the reproductive tract result in increased recombination? Here we answer these questions using classical genetics in *Drosophila melanogaster* with different *Wolbachia* variants and using another intracellular symbiont, *Spiroplasma poulsonii.* We confirm that *Wolbachia* significantly increases recombination across two intervals, one on the X and one on the 2^nd^ chromosome, but we could not detect any effect on the 3^rd^ chromosome interval queried. In addition, there is a clear correlation between *Wolbachia* load and recombination events, suggesting *Wolbachia* itself is the cause of the elevated recombination; clearing the host of *Wolbachia* restores recombination rate to a basal level while infection with a high-titer variant increases recombination. Another intracellular symbiont, *Spiroplasma*, does not increase recombination rate, suggesting this phenomenon is not simply due to the presence of a bacterial infection in the gonads, but is *Wolbachia* specific. These results suggest that *Wolbachia* specifically elevates host recombination, providing a previously unknown benefit to its host.

## Results

### Wolbachia infection increases host recombination rate

We reasoned that if *Wolbachia* infected flies exhibited increased recombination rate, this would be evident in natural populations. We took advantage of a set of isogenized flies, sampled from a wild-caught population in North Carolina, the Drosophila Genetic Reference Panel ^17^. Virgin females from two backgrounds (DGRP-320, infected with *Wolbachia* and DGRP-83, *Wolbachia-free*) were crossed independently to three different lines carrying chromosomal markers, allowing us to distinguish recombinants along certain genomic intervals based on presence of dominant markers (**Fig. 1**). These lines were y[1] v[1] (BDSC stock #1509) for the X chromosome, carrying *yellow* and *vermillion*; vg[1] bw[1] (stock #433) for the 2^nd^ chromosome, carrying *vestigial wings* and *brown*, and e[4] wo[1] ro[1] (stock #496) for the 3^rd^ chromosome, carrying *ebony* and *rough*.

**Figure 1.**
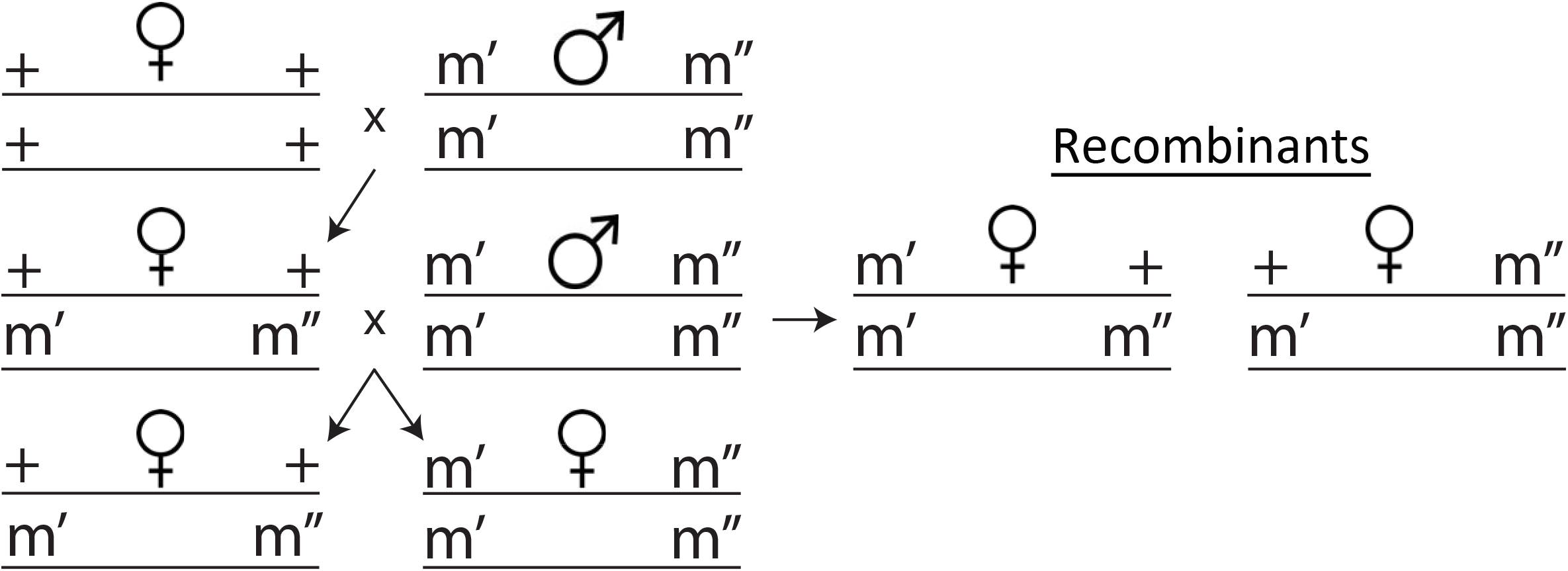
The crossing pattern used to track recombination events. ++ refers to wild type and m’m” refers to genetic markers on each chromosome (*y v* on the X; *vg bw* on the 2nd; *e ro* on the 3rd). Recombination is tracked by looking at the ratio of recombinants to the total number of progeny produced.

As a control, we also cleared the *Wolbachia* infection from line DGRP-320 by rearing the flies on tetracycline for 3 generations and then repopulating the extracellular microbiome for 1 generation. When we compared *Wolbachia* infected and tetracycline cleared individuals, controlling for genetic background, we observed a statistically significant increase in recombination in F2 progeny derived from *Wolbachia-infected* mothers (**Fig. 2**). Specifically, for the X and 2^nd^ chromosomes we observed an increase in mutation rate of 6.4% and 6.1%, respectively. No statistically significant effect of *Wolbachia* infection was observed on the 3^rd^ chromosome (df = 14, ⍰2 = 0.080, p = 0.77). It may be that this difference in recombination rate across chromosomes reflects the natural variation in recombination observed across genomic intervals for *Drosophila*, or may be an artifact of the genomic interval sampled and not an influence of *Wolbachia* on specific chromosomes.

**Figure 2.**
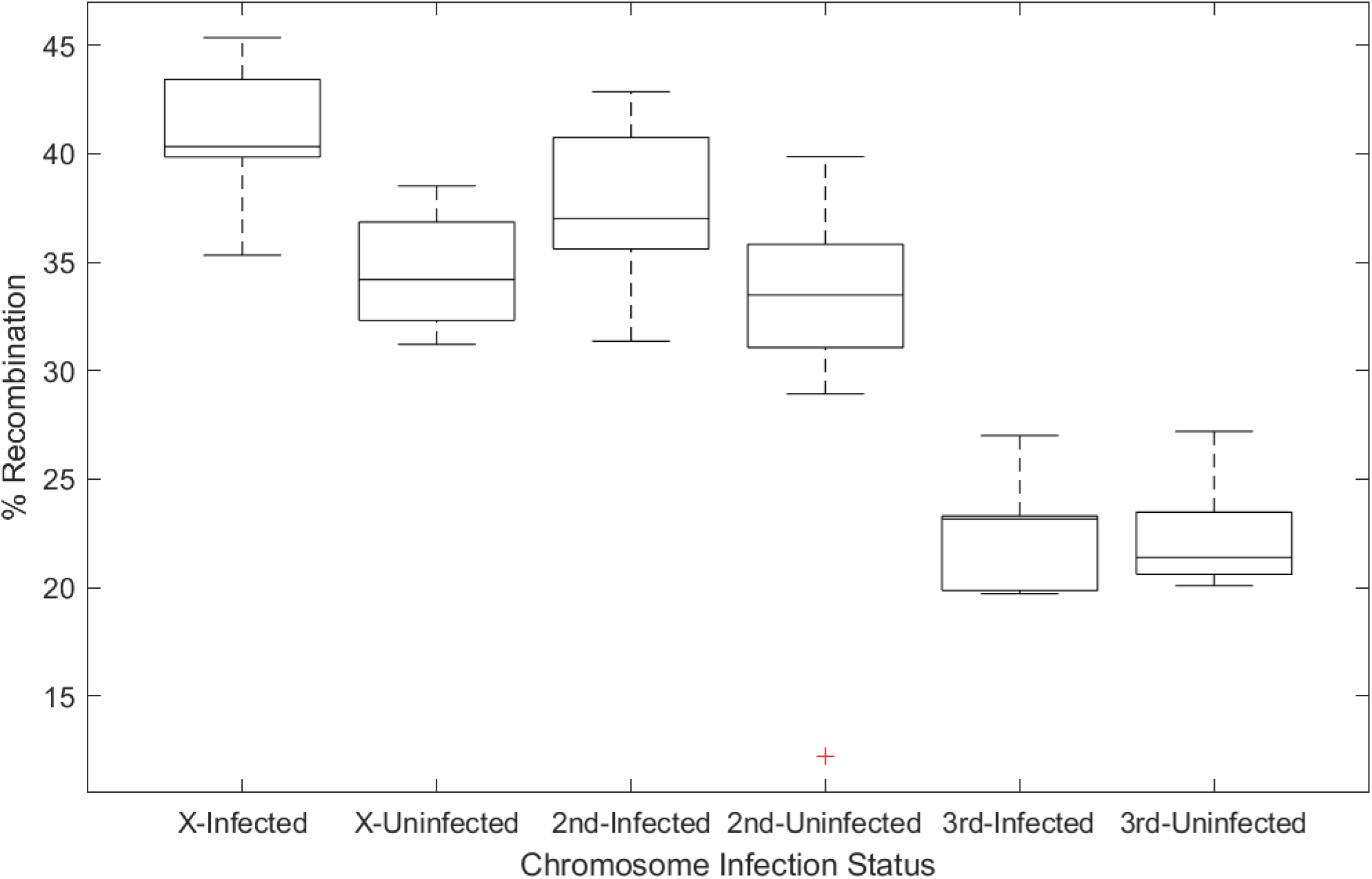
Percentage of recombinants observed, based on genetic markers on the X, 2nd, and 3rd chromosome of *Drosophila*. *Wolbachia* infection significantly increased the recombination rate observed on both the X and 2nd chromosome by 6.4 and 6.1%, respectively. One way ANOVA produced significance for the X (df = 17, χ^2^ = 16.084, p < 6.058e^-5^) and the 2nd chromosome (df = 17, χ^2^ = 7.2357, p < 0.007147), but not on the 3rd chromosome (df = 14, χ^2^ = 0.080081, p = 0.7772).

### Dose dependent effect of Wolbachia on host recombination rate

Because we observed a strong influence of *Wolbachia* infection status on host recombination rate, we sought to modulate infection status by using high titer *Wolbachia* infections in our experiment. *Wolbachia* colonize *Drosophila* at different titers depending on the amplification of a specific genomic interval in the *Wolbachia* genome termed “octomom” ^18^. We crossed females carrying the highest titer, pathogenic *Wolbachia* variant, *w*MelPop, (w[1118]/Dp(1;Y)y[+] |Wolbachia-*w*Melpop; BDSC stock #65284), with males of stock DGRP-320. Lines were introgressed for 3 generations within the #DGRP-320 genetic background before use in experiments. We looked specifically at the X chromosome intervals as we had already established that *Wolbachia* significantly increased recombination across that genomic interval (**Fig. 2**). Again, we observed a significant effect of *Wolbachia* infection on recombination rate in this experiment (one-way ANOVA; df = 15, χ^2^ = 15.14, p = 0.015) (**Fig. 3**). Interestingly, we observed a significant effect of *Wolbachia* titer on recombination rate – the high titer *w*MelPop variant increases recombination on the X chromosome in F2 progeny by 9.5% compared to the 6.3% observed for *w*Mel (df = 11, χ^2^ = 12.65, p = 0.044) (**Fig. 3**).

**Figure 3.**
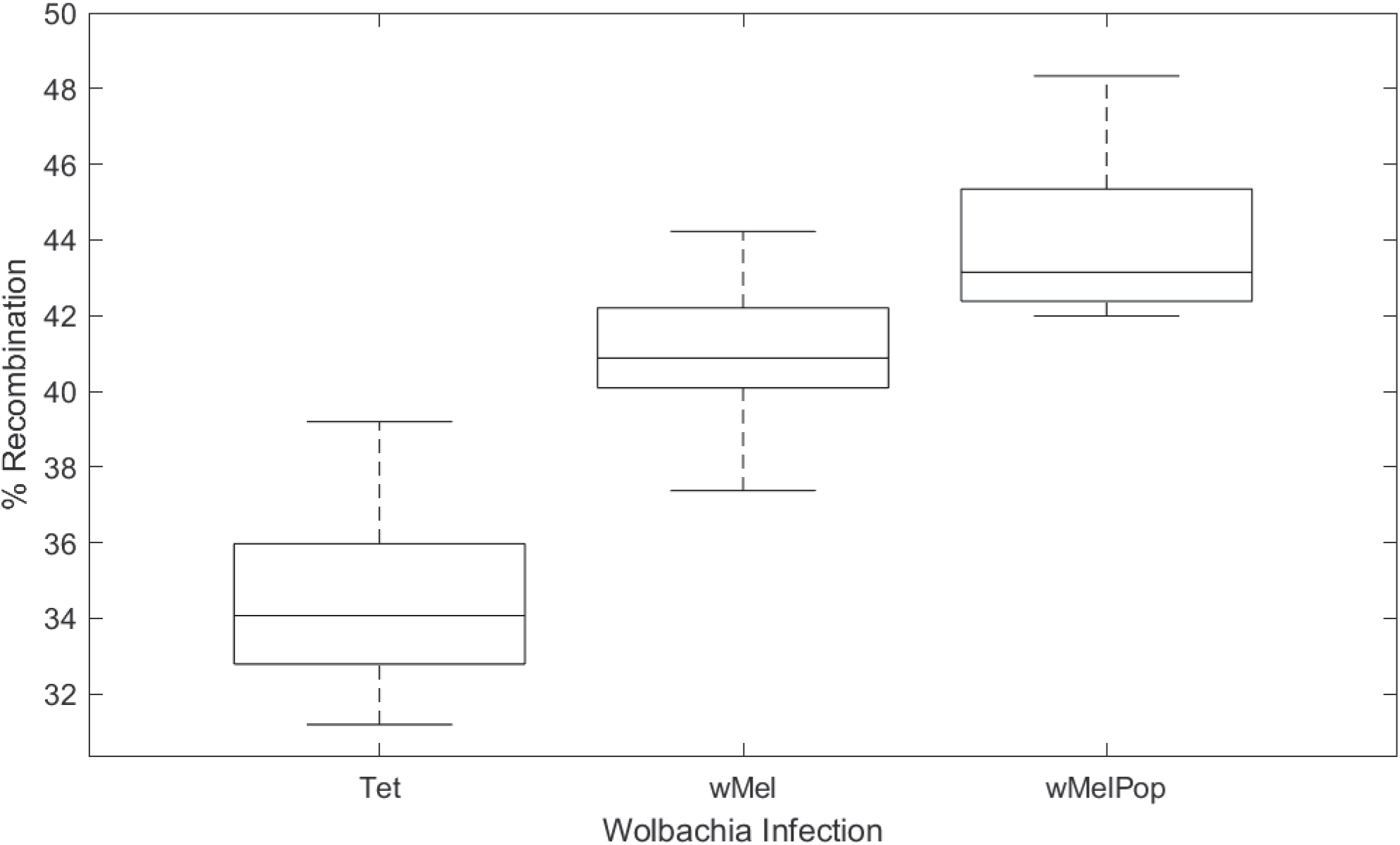
Percentage of recombinants observed, based on genetic markers on the X chromosome of *Drosophila*, when *Wolbachia* titer is varied. A uniform genetic background was used for comparisons across *Wolbachia*-uninfected (Tet) flies, *Wolbachia* infected (*w*Mel) flies, and flies infected with a high-titervariant (*w*MelPop). Increased *Wolbachia* load increased recombination in a load-dependent manner by 6.3% for *w*Mel and 9.5% for *w*MelPop. Significance between titer load was determined by one way ANOVA (df = 15, χ^2^ = 15.1495, p = 0.0153). Significance between Tet and *w*Mel (df = 11, χ^2^ = 15.536, p < 8.094e^-5^), *w*Mel and *w*MelPop (df = 11, χ^2^ = 12.652, p = 0.0442), and Tet and *w*MelPop (df = 11, χ^2^ = 25.994, p < 3.424e^-7^) was determined by one way ANOVA.

### Spiroplasma does not increase host recombination rate

We hypothesized that *Wolbachia* may be a stress on the host cell, increasing recombination rate as a result of increased reactive oxygen species or other immune activation pathways. We therefore reasoned that any bacterial infection may increase recombination rate. To test this hypothesis, we procured *Spiroplasma poulsonii* MSRO (a gift from John Jaenike), which we used to infect a *Wolbachia-free* OreR lab stock (Oregon-R-modENCODE, BDSC #25211). We used the same crossing scheme as above to introduce *Spiroplasma* into the y[1] v[1] background, carrying phenotypic markers on the X chromosome. As a genetic control, we used stock #25211. Counter to our hypothesis, we observed no significant increase in recombination based on *Spiroplasma* infection (**Fig. 4**).

**Figure 4.**
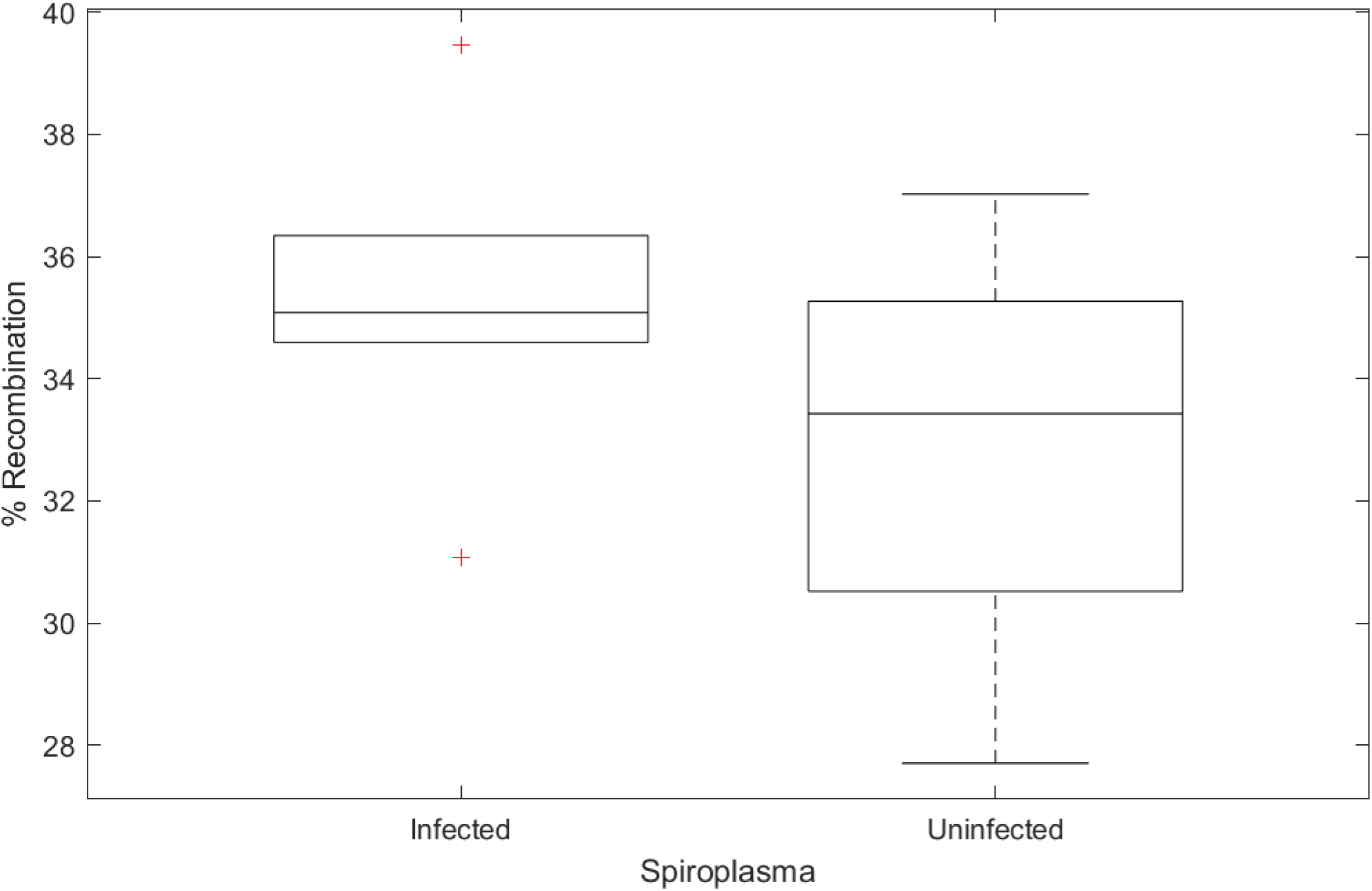
Percentage of recombinants observed based on genetic markers on the X chromosome of *Drosophila. Spiroplasma* infection did not significantly increase the observed recombination rate through one way ANOVA (df = 11, χ^2^ = 0.03885, p = 0.2153).

## Discussion

For sexually reproducing organisms, recombination is both a source of genetic diversity within a population and a mechanism by which to decouple differential selection on sites across the chromosome. Therefore, recombination is thought to be beneficial. Here, we observed that *Wolbachia* infection significantly increased the recombination rate observed across two genomic intervals (for both the X and the 2^nd^ chromosome). Importantly, two lines of evidence presented here support the hypotheses that this increase is *Wolbachia* specific: we observed a dose dependency to the recombination rate and we did not identify a significant effect of *Spiroplasma* infection on recombination rate. Recombination rates vary dramatically across animals, even within a genus, as best illustrated within the *Drosophila* clade^19,20^. The mechanism behind this difference is not well understood but our data suggest that the symbiont *Wolbachia* may influence recombination rate of infected *Drosophila*. This result suggests a previously unknown benefit to *Wolbachia* infection and may help explain the prevalence of *Wolbachia* in certain insect populations.

The mechanism by which *Wolbachia* infection elevates recombination is an active area of inquiry in our lab. *Wolbachia* have an active type IV secretion system that they use to secrete proteins into the host and modulate host cell biology. It is possible that some of these proteins may influence recombination rate directly or indirectly, although no effectors have yet been identified that bind to host DNA. Here we used two different *Wolbachia* variants to support the hypothesis that *Wolbachia* increases host recombination rate. However, it is possible that strains outside of the *w*Mel clade do not increase host recombination and a comparative genomic framework could be used to identify loci in *Wolbachia* that confer the phenotype. Finally, a recent publication suggested *Wolbachia w*Mel infected *Drosophila* prefer cooler temperatures ^21^. Increases in temperature modulate recombination in *Drosophila* ^22^ and it is possible that *Wolbachia* infection elevates host temperature enough to generate an increase in the number of detected recombinants, in a laboratory setting where flies are kept at a constant temperature.

*Wolbachia* is known for its ability to transfer between species. This process can occur through horizontal transmission of the strain or through introgression via hybridization ^23^. One particular strain, *w*Ri, has been identified as having globally spread across highly divergent *Drosophila* species, and in a few cases, instances of introgression between species are known to have facilitated this transfer ^23^. *Wolbachia* facilitates its own maintenance in populations through reproductive manipulations ^5^ and potentially through mutualistic benefits offered to the host ^6,11^. *Wolbachia* has been shown to facilitate divergence of hosts, through manipulation of sperm-egg compatibility, strengthening species boundaries ^24^. It is therefore tempting to suggest, based on these results, that *Wolbachia* may also increase introgression between species to facilitate their own spread. This hypothesis, however, awaits further research.

## Acknowledgements

We would like to thank anonymous reviewers for their feedback. We also thank MaryAnn Martin for their assistance and guidance throughout the course of this project and Nadia Singh for discussions at earlier stages of data analysis. This work was funded in part by the National Science Foundation (NSF 1456545 to ILGN).

## Author contributions

ILGN and KBS conceived of the project, ILGN, and KBS implemented the experiments, and wrote the manuscript.

## Data and Reagent Availability

*Drosophila* strains used in this work are publicly available through the Bloomington Drosophila Stock Center. *Spiroplasma* infected OreR is available from the Newton laboratory – please contact.

## Methods

### Fly Rearing

Flies were ordered from the Bloomington Drosophila Stock Center. Three marker stocks were selected as recombination trackers for the X, 2^nd^, and 3^rd^ chromosomes in *Drosophila melanogaster:* #1509, which is marked with *yellow* (*y*) and *vermillion* (*v*) on the X chromosome; #433, which is marked with *vestigial wings* (*vg*) and *brown* (*bw*) on the 2^nd^ chromosome; and #496, which is marked with *ebony* (*e*) and *rough* (*ro*) on the 3^rd^ chromosome. Two stocks, one *Wolbachia* infected and one uninfected, were selected at random from the Drosophila Genetic Reference Panel (DGRP): #29654 and #28134, respectively. To modulate infection status, we introduced high titer *Wolbachia* infections into our stock #29654 using the *Wolbachia* variant *w*MelPop from stock #65284. To clear #29654 of its infection, flies were raised on fly food containing 50 ug/mL tetracycline for 3 generations and the allowed to be recolonized by their extracellular microbiome, and recover from tetracycline, for 1 generation. All crosses were conducted at room temperature.

### DNA Extraction and Polymerase Chain Reaction

*Wolbachia* infection in #29654 was confirmed by PCR. DNA was extracted using a single-fly extraction method. Whole flies were ground with a pipette tip containing 50 microliters lysis buffer (10mM Tris-HCl pH 8.2, 1 mM EDTA, and 25 mM NaCl) and 5 microliters Proteinase K. They were incubated at room temperature for 20 minutes then heated to 95°C for 2 minutes to deactivate the enzyme. Polymerase chain reaction was performed on standardized quantities of extracted DNA. The cycling conditions are as follows: 98°C for 2 minutes, followed by 30 cycles of 98°C for 30 seconds, 59°C for 45 seconds, and 72°C for 1 minute 30 seconds, then finished with 72°C for 10 minutes. Primers used for this are as follows: wsp F1 5’-GTC CAA TAR STG ATG ARG AAA C -3’ and wsp R1 5’-CYG CAC CAA YAG YRC TRT AAA -3’. Amplified *Wolbachia* DNA was visualized using agarose gel electrophoresis. Quantitative PCR was performed to confirm the titer difference in *Wolbachia* infection between *w*Mel to *w*MelPop. Data were collected using an Applied Biosystems StepOne Real-time PCR system and iTaq universal SYBR Green supermix. The *Wolbachia* primers used are as follows: wspF 5’-CATTGGTGTTGGTGTTGGTG -3’ and wspR 5’-ACCGAAATAACGAGCTCCAG -3’. The host primers used are as follows: Rpl32F 5’-CCGCTTCAAGGGACAGTATC -3’ and Rpl32R 5’-CAATCTCCTTGCGCTTCTTG -3’. The cycling conditions are as follows: 50°C for 2 minutes, 95°C for 10 minutes, followed by 40 cycles of 95°C for 30 seconds and 59°C for 1 minute. The reaction was carried out in a 96-well plate. Gene expression was determined by the Livak and Pfaffl methods.

### Recombination Assay

To determine if recombination events had occurred, a two-step crossing method was devised, shown in Figure 1. Ten virgin DGRP females aged 3-5 days were housed with ten phenotypically marked males and were allowed to mate for 10 days, after which parentals were cleared from the bottle. Virgin female F1 progeny were collected and crossed to the male parental line in the same ratio as before and allowed to mate for 10 days before being cleared from the bottle. All F2 progeny from this cross were collected and frozen after 10 days of the clearing. Flies were scored according to their visible phenotypes and sorted into two groups. Significance between phenotypically normal flies and flies that have undergone recombination was determined through ANOVA.

